# Multi-omics integration and batch correction using a modality-agnostic deep learning framework

**DOI:** 10.1101/2025.10.21.683449

**Authors:** Jose Ignacio Alvira Larizgoitia, Gabriele Partel, Lorenzo Venturelli, Wanqiu Zhang, Xander Spotbeen, Sebastiaan Vanuytven, Sam Kint, Katy Vandereyken, David Wouters, Anis Ismail, Regis Scarceriaux, Jakub Idkowiak, Tassiani Sarretto, Shane R. Ellis, Massimo Loda, Fabio Socciarelli, Thomas Gevaert, Steven Joniau, Marc Claesen, Nico Verbeeck, Thierry Voet, Johannes V. Swinnen, Jelle Jacobs, Alejandro Sifrim

## Abstract

State-of-the-art biotechnologies allow the detection of different molecular species on the same biological sample, generating complex highly-dimensional multi-modal datasets. Gaining a holistic understanding of biological phenomena, such as oncogenesis or aging, requires integrating these diverse modalities into low-dimensional data representations while correcting for technical artifacts. Here we present MIMA, a modular, unsupervised AI framework for multi-omics data integration and batch correction. Applied to complex spatial and single-cell datasets, MIMA effectively removes batch effects, while preserving biologically relevant information, and learns representations predictive of expert pathologist annotations. Additionally, it enables cross-modal translation, uncovers molecular patterns not captured by manual annotations, and despite being modality-agnostic performs on par with specialized state-of-the-art tools. MIMA’s flexibility and scalability make it a powerful tool for multimodal data analysis. MIMA provides a foundation for AI-based, multi-omics augmented digital pathology frameworks, offering new opportunities for improved patient stratification and precision medicine through the comprehensive integration of high-dimensional molecular data and histopathological imaging.

## Introduction

Characterizing the complex biology of heterogeneous tissues requires a comprehensive understanding of the molecular profiles and interactions that define diverse cellular states. To achieve this, contemporary research increasingly relies on the integration of multiple molecular layers, such as transcriptomics, genomics, proteomics and metabolomics, each offering complementary insights into cellular function and identity^1^. These datasets can uncover fine-grained cellular and microenvironmental heterogeneity and provide insights into cross-modal regulatory mechanisms^2,3^. The data modalities differ however substantially in their dimensionality, noise characteristics, and scale, and are often confounded by large batch effects introduced by variations in sample preparation, instrumentation, and experimental protocols. These challenges hinder robust multisample, cross-modal analysis and limit our ability to extract biologically meaningful insights from multimodal datasets.

Combining heterogeneous data inputs into unified representations, while correcting for batch and other modality-specific technical confounding effects, requires computational data integration frameworks^4^. The challenge of multi-omics data integration is complicated by inherent differences in the experimental data generation processes, leading to distinct distributions (i.e. gene expression counts compared to ion intensities in mass spectrometry) and variance components, which can be either technical or biological in nature. This is especially problematic where multi-modal data is collected from different biological replicates, adjacent tissue sections or across different technology platforms for the same modality. Unaddressed, these artifacts obscure true biological signals and may undermine downstream analyses^5^. Therefore, a crucial step towards robust multimodal integration is the ability to disentangle distinct sources of variation. In the context of single-cell omics, this means explicitly separating shared variability across modalities from modality-specific, batch-related and potentially other covariates^6^. Without this disentanglement, integrated representations risk conflating meaningful biology with measurement artifacts, reducing interpretability and accuracy. Properly modelling these differences may not only improve self-supervision of the models but also support biologically grounded interpretation of the learned representations, cross-modal translation, and more accurate downstream tasks like clustering, trajectory inference, and classification^1,7,8^.

Several computational approaches have been developed to address various aspects of the data integration problem. However, many are specifically designed for the integration of particular pairs of modalities (for example *multiVI*^*9*^ or *totalVI*^*10*^) and often lack the flexibility to be readily extended to incorporate additional omics layers. Linear factorization models like *MOFA+*^11^ and graph-based approaches like *GLUE*^*12*^ extend to three modalities, but often assume that all samples are profiled with the same set of modalities or that the features (e.g., genes, proteins) are directly comparable across datasets. Newer tools such as *scVAEIT*^*13*^, *scMoMaT*^*14*^, *StabMap*^*15*^, *Multigrate*^*16*^, and *MIDAS*^*17*^ integrate ‘mosaic’ datasets, where some modalities are missing per sample, but typically lack the ability to simultaneously perform batch correction or disentangle biological signals.

To address these limitations, we present MIMA (Multimodal Integration with Modality-agnostic Autoencoders), a scalable and modality-agnostic computational framework using representation learning for the integration and batch correction of paired multimodal datasets. MIMA explicitly disentangles shared biological signals, modality-specific variation, and batch-related technical artifacts through separate latent spaces. This design enables both robust data harmonization and biologically meaningful integration. Unlike many domain-specific models, MIMA is self-supervised and modality-agnostic, requiring no feature selection or specialized preprocessing. It supports paired multimodal datasets and enables accurate cross-modal translation by learning a shared representation from which missing modalities can be predicted.

Specifically, MIMA’s modular architecture allows each modality to be modelled using tailored encoders and decoders, while jointly contributing to a product-of-experts latent space^18^ that captures shared biology. Its private latent spaces retain modality-specific features that would otherwise be lost or misinterpreted in a purely shared representation. This structure enhances interpretability, improves reconstruction quality, and allows the model to scale across different data types and biological contexts.

We showcase MIMA’s modular and modality agnostic architecture by integrating three modalities (i.e. Visium-based spatial transcriptomics, Mass Spectrometry Imaging-based lipidomics and H&E derived morphological features). While gene expression and MSI measurements have been integrated before using linear methods^19,20^, no AI-based architecture exists that can capture the expected non-linearities and generate common embeddings between these modalities. MIMA effectively removes batch effects, enables cross-modal translation, and learns representations highly predictive of domain-expert manual annotations. We further validate the framework using two additional multimodal benchmarks: a 10x multiome dataset combining gene expression and chromatin accessibility, and a CITE-seq dataset pairing transcriptomic and protein abundance profiles. MIMA’s benchmarking performance matches or exceeds that of specialized tools while offering broader applicability. Its generalizable design and interpretability make it a powerful framework for multimodal omics, and a promising foundation for future AI-augmented digital pathology systems.

In summary, the proposed MIMA framework enables the disentangled representation of multimodal biological data by explicitly modelling shared biological signals, modality-specific information, and batch effects in separate latent spaces. Its modular architecture supports diverse data types and facilitates both integration and batch correction through a unified approach. By combining modality-specific encoders, a shared Product-of-Experts representation, and separate loss functions, the model produces biologically meaningful, denoised, and batch-corrected reconstructions, as well as accurate cross-modal translations. This makes it a powerful and generalizable tool for robust analysis of complex, paired multi-omics datasets.

## Results

### The MIMA model

MIMA (Multimodal Integration with Modality-agnostic Autoencoders) is a scalable and modality-agnostic deep learning framework for the integration and batch correction of paired multimodal datasets. It is designed under the assumption that multimodal biological data comprises multiple, distinct sources of variability. These include: (i) shared biological variability, which reflects biological signals that are present across modalities; (ii) modality-specific biological variability, representing biologically meaningful variation that is uniquely captured within each modality; and (iii) technical variability, specific to each modality and arising from differences in experimental conditions (i.e. reagent lots, experimental runs, sample handling) commonly referred to as *batch effects*. For example, shared biological variability may involve changes in the expression of genes implicated in lipid metabolism and their corresponding effect on the abundance of specific lipid species. Conversely, modality-specific variability includes gene expression patterns which are uncorrelated with changes in lipid profiles, but are nonetheless informative for understanding cellular heterogeneity.

Based on these assumptions, we propose a modular multimodal variational autoencoder (MVAE) architecture (Fig. 1A), comprising separate encoder–decoder submodules for each data modality. This design allows flexibility in adapting to diverse data types such as tabular, image, or sequence data. Each modality-specific encoder maps the input into three distinct latent representations: (i) a shared latent space, intended to capture biological signals common across all modalities; (ii) a modality-specific latent space, which encodes the biological information captured by that modality; and (iii) a batch latent space, designed to isolate variability unique to each sample or batch.

**Figure 1.**
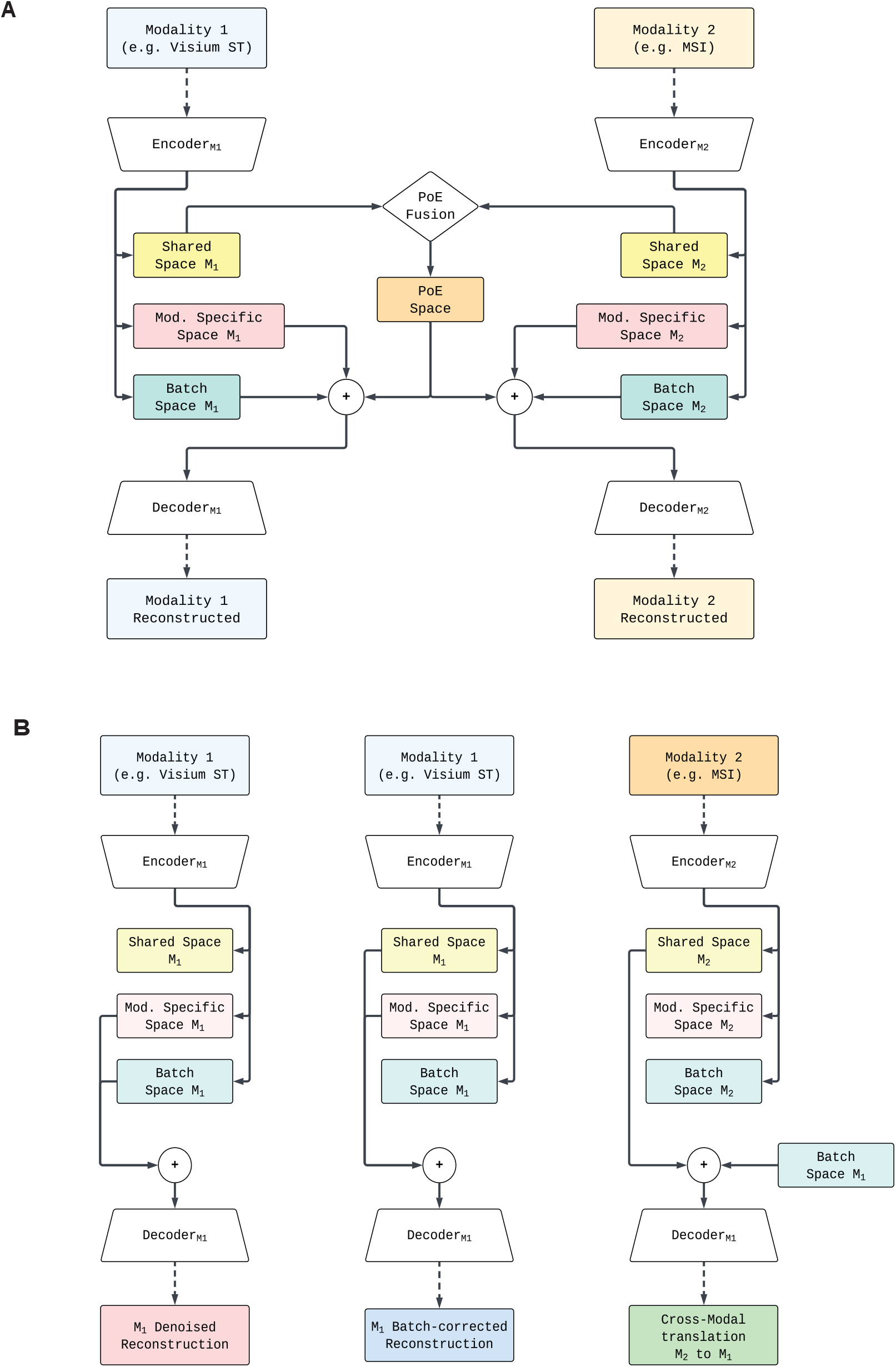
MIMA model architecture. (A) Schematic of MIMA’s architecture. Each modality has its own encoder-decoder pair, projecting into shared and modality-specific latent spaces, combining the shared latent spaces via a product-of-experts. (B) MIMA’s outputs produced by combining different latent spaces.

To account for batch effects, MIMA models a separate latent space for each sample, ensuring that sample-specific variation is isolated (Methods). This architectural constraint promotes the disentanglement of sample-specific variation most likely representing technical noise or sample-specific artifacts and biology. To construct a unified representation across modalities, the shared latent spaces are combined using a Product-of-Experts (PoE) formulation^18^, ensuring that only information consistently present in all modalities is retained. The modality-specific latent spaces are left unaltered, as they are designed to capture unimodal biological variability. The final latent representations, comprising the shared, batch, and modality-specific components, are then summed up and passed through their respective decoders to reconstruct the original input data. This architecture enables three types of outputs for each modality: (i) denoised reconstruction, by decoding the combination of shared, modality-specific, and batch latent spaces; (ii) cross-modal translation, by decoding the shared latent space in combination with the batch latent space from a target modality; and (iii) batch-corrected reconstructions, by decoding the shared and modality-specific latent spaces, excluding batch information (Fig. 1B).

To specifically model these different sources of variation, the model is optimized using a composite loss function that separates shared and private objectives, extending the idea presented in Lee and Pavlovic^21^ (Methods). The shared loss is computed as the mean squared error (MSE) between the input data and both (i) the reconstruction generated from the PoE-shared latent space, and (ii) the cross-modal translation output. The latter encourages accurate representation of shared biological information without influencing the batch or modality-specific latent spaces. A Kullback-Leibler (KL) divergence regularization term is applied to the shared latent space using a standard normal prior. The private loss accounts for the reconstruction error from the batch and modality-specific latent spaces. Here, the MSE is computed between the original input and its reconstruction using only the private components. Separate KL divergence terms are applied to the batch and modality-specific latent spaces to similarly constrain them to standard normal distributions.

### MIMA corrects batch effects and integrates spatial transcriptomics, spatial lipidomics and tissue morphology of a prostate cancer dataset

To illustrate the functionality of our framework, we applied MIMA to a novel prostate cancer spatial multi-omics dataset, consisting of 10X Genomics Visium spatial transcriptomics, MSI lipidomics and histological imaging^19^ (Fig. 2A, Methods). UMAP-based dimensionality reduction of the input data shows that all 3 modalities have significant inter-sample differences, i.e. batch effects, which are common for these spatial omics technologies^22–24^ and can occlude underlying biological signals (Fig. 2A). This was most pronounced in the MSI data, where batch effects can hamper multi-sample data integration^23^.

**Figure 2.**
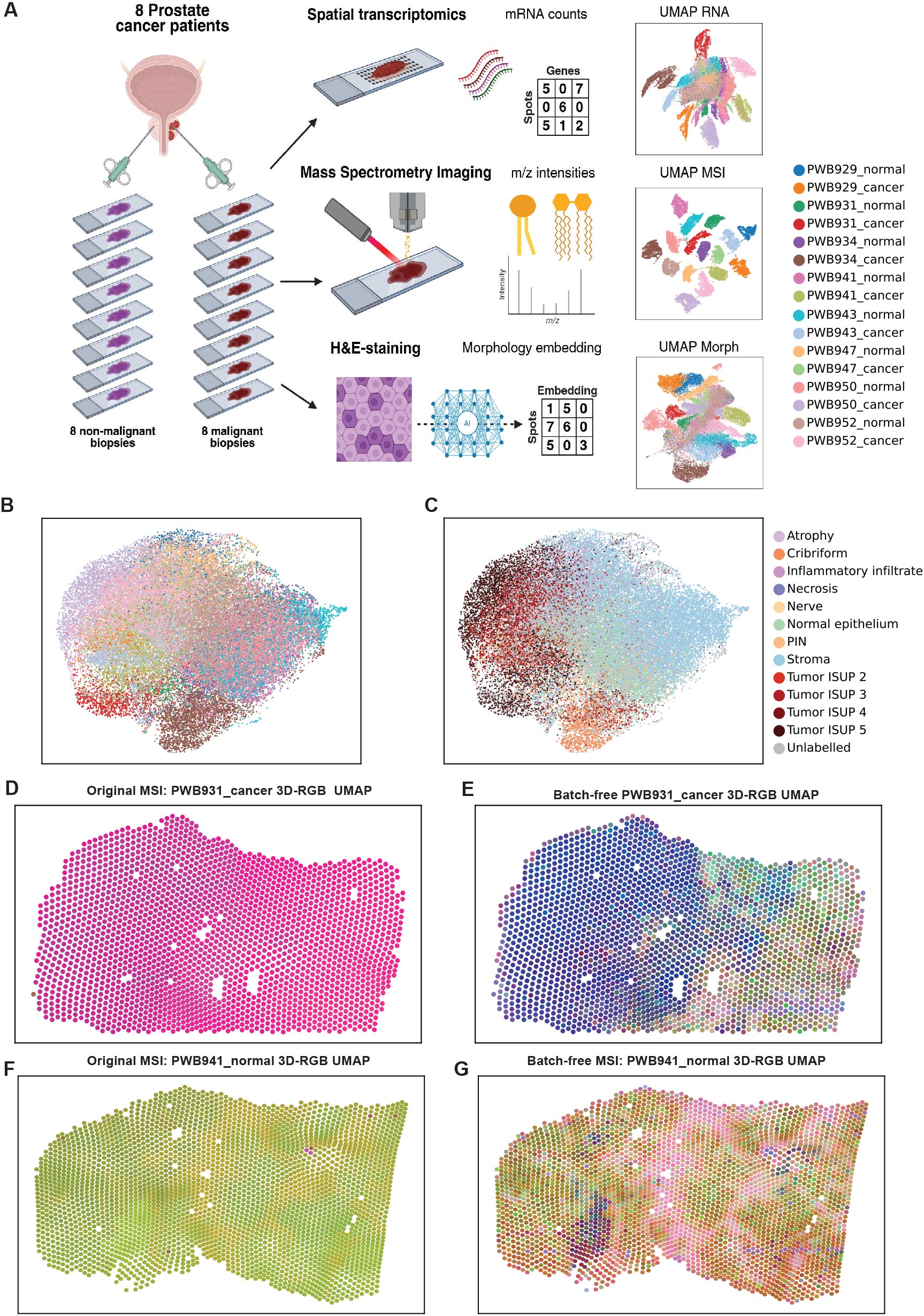
Prostate cancer dataset and batch correction performance of MIMA. (A) Overview of the primary prostate cancer dataset used in this study. Two tissue sections were taken from each of the eight patients: one tumor-proximal and one tumor-distal, each profiled using Visium spatial transcriptomics, mass spectrometry imaging (MSI), and H&E-based morphology embeddings. UMAP visualizations of the input data for all three modalities. (B-C) UMAP embeddings of MIMA’s batch-corrected data for the MSI modality, colored by sample (B) and by pathologist annotations (C), showing alignment across samples after batch correction. To visually interpret the structure of the learned latent space, we compute a 3D UMAP embedding from the latent representations and map each spot to an RGB color, this encoding allows each spatial spot to be assigned a unique color based on its position in the 3D embedding space. (D-E) 3D-RGB UMAP visualizations mapped onto the tissue section for sample PWB931_cancer for input, uncorrected, MSI data (D) and MIMA’s batch-corrected output (E). (F-G) 3D-RGB UMAP visualizations mapped onto the tissue section for sample PWB941_normal for input, uncorrected, MSI data (F) and MIMA’s batch-corrected output (G).

MIMA produces batch-corrected outputs by reconstructing the data using only the shared (PoE) and modality-specific latent spaces, excluding the batch latent space to suppress technical variability (Fig. 2B-C and Supplementary Fig. 2A and 3A). The corrected representations demonstrate improved alignment between samples and reveal structure consistent with domain-expert manual annotations. Before batch correction, we failed to observe meaningful spatial molecular patterns in the MSI data, since every sample is its own cluster (Fig. 2D & F). This prevents multisample comparative analysis, a known issue in the MSI field^23^. The MIMA batch-corrected outputs reveal clear spatial patterns and capture multi-sample biological signals (Fig. 2E & G), indicating that MIMA successfully disentangles and removes batch effects, while preserving biologically relevant variability. Compared to the lipidomics modality, the transcriptomics and histology modalities exhibited less pronounced batch effects (Fig. 2A, UMAP RNA and Morphology). Nonetheless, in both modalities, certain samples still formed distinct clusters in the raw data, indicating residual technical variation. After processing the RNA data through the trained MIMA, the batch-specific clustering was largely removed, while the underlying biological structure was preserved (Supplementary Fig. 2A). Notably, benign-like annotated regions remained distinct from malignant or tumor-associated regions, supporting the model’s ability to retain biologically meaningful variation. Spatial visualization of the batch-corrected transcriptomics data (Supplementary Fig. 2 B-E) revealed clear spatial patterns at the tissue-level. The integrated output showed more refined, higher-resolution biological gradients across samples, further validating the utility of the MIMA in enabling cross-sample analyses.

In the histology modality, batch correction yielded more modest improvements in biological clustering, likely due to the lower dimensionality of the input features (512D) and the fact that this modality was derived from H&E-stained images using a pre-trained deep learning model^25^. Nevertheless, the spatial representation of the batch-corrected morphological embeddings (Supplementary Fig. 3 B-E) exhibited higher detail granularity compared to the uncorrected embeddings. These results underscore MIMA’s capacity to recover biologically informative signals across heterogeneous datasets that are typically hindered by significant technical variability.

### Combining biological information in a shared latent representation

Besides automated batch correction, our model also learns relationships between the different data modalities, enabling the translation between modalities and to generate shared latent representations that effectively integrate multi-modal biological signals. To evaluate the integrative capabilities of our framework and its ability to extract biologically meaningful features, we inspected both the shared latent representation (PoE space) and the modality translation outputs on the prostate cancer (PCa) dataset. Specifically, we evaluated MIMA by assessing the structure of the integrated latent space and its predictive performance on domain-expert manual annotations.

Qualitatively, the PoE latent space exhibits separation of manual annotations, indicating a well-integrated representation (Fig. 3A). Data points annotated by the pathologist as benign-like or malignant remain distinct. Three samples (PWB929_cancer, PWB931_cancer, and PWB934_cancer) appear as outliers in the integrated space. These samples contain rare annotation classes (Supplementary Fig. 1C), making their integration more challenging. However, the preservation of these distinct clusters suggests that MIMA avoids overcorrecting sample-specific variability as batch effects and instead retains rare, potentially meaningful biological variation in the shared latent space.

**Figure 3.**
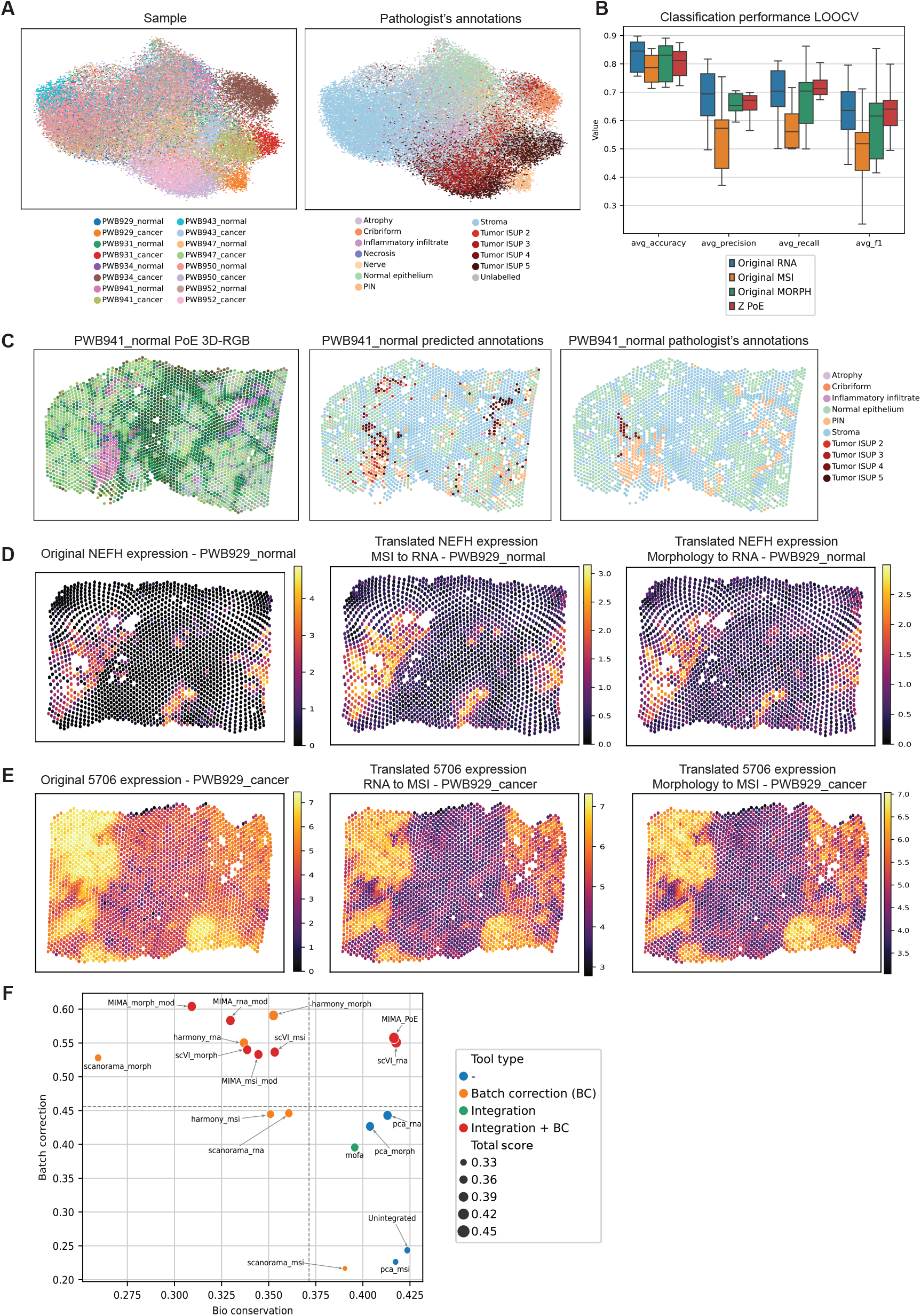
Integration and translation across RNA, MSI, and morphology. (A) UMAP of the PoE latent space, colored by batch (left) and pathologist annotations (right), demonstrating modality integration, batch correction and biological structure preservation. (B) Leave-one-out cross-validation (LOOCV) classification performance for each input modality individually and for the integrated PoE latent space, evaluated using average accuracy, precision, recall, and F1 score. To visually interpret the structure of the learned latent space, we compute a 3D UMAP embedding from the latent representations and map each spot to an RGB color, this encoding allows each spatial spot to be assigned a unique color based on its position in the 3D embedding space. (C) Left: 3D-RGB UMAP of the PoE latent space, highlighting learned spatial patterns. Middle: predicted pathology annotations using 4-fold CV ensemble classifier (EC). Right: ground-truth pathologist annotations. (D) Cross-modal gene expression reconstruction: predicted NEFH expression from MSI and morphology compared to the actual spatial transcriptomics data. (E) Cross-modal lipid reconstruction: predicted abundance of lipid 5706 from RNA and morphology, compared to the true MSI measurement. (F) Quantitative benchmarking of MIMA and alternative methods. X-axis shows biological conservation, Y-axis shows batch correction performance. Each point represents a method, with dashed lines indicating the overall averages.

To quantitatively assess the biological information encoded in the PoE space, we trained an ensemble of random forest classifiers (EC, Methods) using the 100-dimensional PoE embeddings as input to predict the 8 biologically relevant annotation classes. We employed leave-one-sample-out crossvalidation (LOOCV) across the 16 PCa samples, training on all-except-one samples and testing on the held-out sample in each iteration (Methods). We report average accuracy, precision, recall, and F1-score across all samples (Fig. 3B). For comparison, the same classification approach was applied to the original, uncorrected transcriptomics, lipidomics, and histology input data. As expected, the classifier trained on lipidomics data performed the worst (average F1 score of 0.49), reflecting strong batch effects encountered in this modality. Transcriptomics (average F1 of 0.57) and histology (average F1 of 0.58) yielded better performance but still performed substantially worse than the combined PoE space (average F1 score of 0.61). Notably, the PoE-based classifier achieved the highest F1-score and exhibited the lowest variance across folds, highlighting the robustness and discriminative power of the integrated representation produced by MIMA.

To explore the spatial distributions of the learned representations, we visualized the PoE embeddings and EC predictions according to their spatial coordinates (Fig. 3C). The integrated latent space captures continuous, fine-grained biological gradients across the tissue, surpassing the granularity of manual annotations. The EC predictions closely align with the pathologist’s labels (average F1=0.63), confirming the predictive power of the PoE space. Remarkably, the model also identifies regions with molecular profiles consistent with tumorigenic states that were not annotated as such by the pathologist. In sample PWB941_normal, classified as benign aside from a small ISUP 4 region, the EC predicts additional tumorous areas within or near zones labelled as Prostatic Intraepithelial Neoplasia (PIN) (Fig. 3C). PIN is a known precancerous condition^26^, and enrichment analysis (Methods: gene enrichment score) of 578 prostate cancer–associated genes (identified by Khan et al.^27^) confirmed that these EC-predicted regions exhibit tumor-like molecular signatures missed by manual inspection (Supplementary Fig. 4A).

Together, these results demonstrate that MIMA not only corrects for technical confounders but also learns biologically meaningful features that enable improved classification and reveal subtle, potentially early-stage pathological changes through the integration of multi-omics and histology.

### Employing the shared latent spaces to translate between different modalities

Imputing missing modalities is highly relevant, especially considering the cost and technical aspects of these experiments. We evaluated the cross-modal translation capabilities of MIMA by testing whether the model could accurately predict missing modalities from observed ones. Translation is achieved by projecting the input modality into the shared latent space, combining it with the batch latent space of the target modality, and decoding it through the corresponding decoder. This approach not only allows the comparison of the imputed translation with the original data (i.e. including that modality’s sample-specific batch effect), but also to its theoretical batch-corrected equivalent.

To qualitatively illustrate this translation process, we selected two representative cases with prior biological information. We examined how well the expression pattern of NEFH, a gene previously associated with prostate cancer^28^, is predicted in sample PWB929_normal using the lipidomic or histology modality as input (Fig. 3D). Using a within-sample four-fold cross-validation (4CV) scheme (where 25% of spatial spots were left out per fold), we reconstructed the entire sample exclusively from test predictions. The translated gene expression maps captured the original spatial pattern of NEFH with similar dynamic range and structure (Pearson’s R^2^ of 0.54 for translation from MSI to RNA and 0.49 from Morphology to RNA), demonstrating that MIMA effectively learns cross-modal relationships and can infer transcriptomic features from metabolic or morphological profiles.

For the lipidomics modality, we visualized translation performance for the m/z value 702.544, associated with lipids PE O-34:2 / PE P-34:1 [M+H]+ which are markers of Tumor ISUP 3 regions in sample PWB929_cancer (Fig. 3E). The transcriptomic and histology modalities were used as input to predict lipidomic features using the previously mentioned cross-validation scheme. The reconstructed lipid maps preserved the spatial enrichment seen in the original MSI data (Pearson’s R2 of 0.77 and 0.76 from transcriptomics or histology respectively), confirming the model’s ability to recover metabolomic signals from transcriptomic or morphological input. Translation to histology was not visualized, as its features are 512-dimensional abstract embeddings derived from H&E stainings. Importantly, this suggests that, after training on large datasets, MIMA could potentially infer multi-omic molecular profiles from standard histopathology images alone, eliminating the need for additional molecular assays.

To systematically quantify the model’s cross-modal translation performance, we evaluated both reconstruction and translation fidelity using a four-fold cross-validation (4CV) scheme (Methods: Classification) ensuring that all evaluated predictions have been obtained from data points never seen by the model. We first computed the mean squared error (MSE) between predicted and observed profiles across all modality pairs. To determine whether MIMA captures spot-specific variation, rather than just reconstructing average cluster-level profiles, we computed the MSE between the input data and the centroid feature vector for that spot’s annotation (i.e. the mean of all other spots carrying the same label). We also contrasted this to the MSE relative to the average feature vector of all spots with different annotations (i.e. all annotations excluding the spot’s own annotation). A lower MSE relative to spots of the same annotation compared to the MSE relative to other annotations indicates that our approach is learning annotation-specific representations, rather than simply capturing global or dataset-wide structures. Specifically, we compared: (i) translation MSE (cross-modal prediction), (ii) reconstruction MSE (same input/output modality), (iii) same-annotation reconstruction MSE, and (iv) different-annotation reconstruction MSE (Supplementary Fig. 4B). To mitigate the effects of sparsity and dropouts for the transcriptomics and lipidomics modalities (Supplementary Fig. 1A), we restricted this analysis to the top 1000 most variable features (Methods: Highly Variable Feature selection, Supplementary Fig. 1B).

Translation and reconstruction MSEs were highly similar across all three modalities, with reconstruction performing slightly better, as expected given the inherent ease of predicting within the same modality. Notably, both reconstruction and translation showed significantly higher fidelity than if the model would have predicted a datapoint belonging to a different annotation. Moreover, cross-modal translation was even of higher fidelity than if the prediction would have been the average profile of spots of that same annotation, indicating that the model captures spot-specific variation rather than simply learning annotation-level averages.

To further assess fidelity and the preservation of feature-wise relationships, we computed Pearson correlation coefficients (R^2^) between the input data and MIMA’s output, under the same 4CV scheme (Supplementary Fig. 4C). These correlations complement the MSE analysis by capturing how well global data structure is preserved in a scale-invariant way. The transcriptomics modality exhibited the broadest R^2^ distribution with the lowest median correlation values, reflecting the inherent sparsity and high dropout rate of transcriptomic data. Nevertheless, the distribution remained skewed towards moderate-to-high correlations, suggesting that the model effectively captures meaningful gene expression patterns in relevant genes. The lipidomics modality showed the highest correlations overall, with narrow distributions around R^2^ ≈ 0.75 – 0.8 for both reconstruction and translation, highlighting the model’s ability to accurately preserve lipid abundance structures. Morphology showed intermediate behavior: although its correlation distributions were broader than MSI’s, they remained significantly above noise level and consistent across both reconstruction and translation outputs. As expected, translation R^2^ values were slightly lower than their reconstruction counterparts for all modalities, reflecting the added difficulty of cross-modal inference. Together with the MSE analyses, these results indicate that MIMA captures feature-level information across all modalities, beyond merely memorizing class averages or annotation identity.

### Benchmarking the bio-conservation and batch correction performance of MIMA

To comprehensively assess the performance of MIMA and compare it with other publicly available tools, we benchmarked its output using the *scIB* metrics package^29^. This commonly used package evaluates integration quality across two categories: bio-conservation metrics, reflecting how well biological annotations are preserved in the embeddings, and batch correction metrics, measuring the degree of batch mixing in the integrated space (Fig. 3F, Methods). We compared our method against tools focused on batch correction (*Harmony*^30^, *Scanorama*^31^, *scVI*^32^), data integration (*MOFA+*^11^), and those that address both integration and batch correction (MIMA). We also included PCA as a baseline, given its widespread use in multi-omics pipelines and as a preprocessing step for models like *SpatialGlue*^33^.

Due to the strong correlation between batch labels and biological annotations in this dataset, a general trend emerges: methods that do not perform batch correction achieve high bio-conservation scores but poor batch correction performance (lower right quadrant), while methods that overcorrect the batch effect show the opposite trend, compromising biological signals (upper left quadrant) (Fig. 3F).

In this context, the PoE space generated by MIMA achieves the highest average score across all metrics, indicating its ability to preserve biological variation while correcting for batch effects. Notably, it is the only tool in the comparison that performs both integration and batch correction effectively. In contrast, *MOFA+*, while effective for integration, is based on an underlying linear model and struggles to resolve complex confounders like batch effects. *scVI*, though not specifically tailored for spatial transcriptomics, lipidomics, or histology, performs reasonably well across modalities, demonstrating its flexibility, even though it cannot produce integrated embeddings. Finally, while the modality-specific latent spaces of MIMA achieve high overall scores, they perform slightly worse in terms of bio-conservation. This suggests that the PoE space successfully integrates and concentrates the most relevant biological information across modalities, making it the most effective representation for downstream analysis.

Moreover, at the time of writing, MIMA is the only tool capable of integrating Visium-based transcriptomics, mass spectrometry imaging (MSI), and histology into a unified framework. In summary, we demonstrate that MIMA not only performs effective batch correction but also learns integrated, biologically meaningful representations across transcriptomics, metabolomics, and histology. These representations enable accurate cross-modal translation, robust biological classification, and uncover spatial disease patterns that go beyond morphology-based annotations. The model is on par with specialized tools for both integration and batch correction, and uniquely supports full multimodal integration of these technologies, positioning it as a powerful framework for comprehensive spatial multi-omics analysis.

### MIMA can be broadly applied to other paired multi-omic datasets

The MIMA framework is designed as a flexible and modular approach that can scale to any number and type of omic input modalities. To demonstrate the generalizability of the model, we applied it to two publicly available single-cell paired multi-omic datasets taken from the 2021 NeurIPS OpenProblems challenge^34^: one combining scRNA-seq with scATAC-seq (10X Genomics multiome), and another combining RNA and protein expression (scCITE-seq^35^) (Fig. 4, Methods). These represent widely used multi-omic datasets with very different modalities compared to lipidomics and histology, offering a valuable benchmark to test the versatility and robustness of our method.

**Figure 4.**
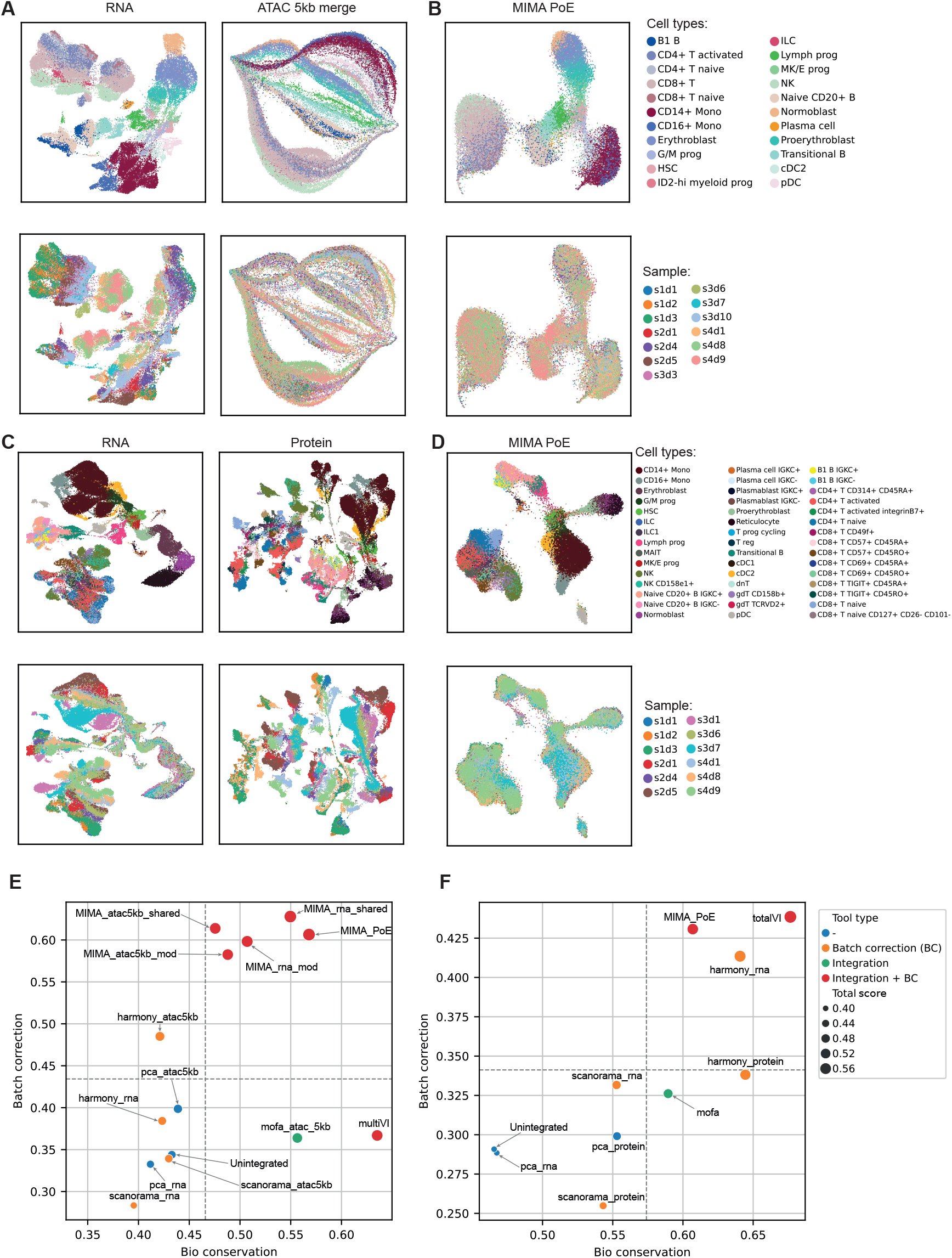
MIMA generalization to multiome and CITE-seq datasets. (A) UMAPs of the preprocessed RNA (left) and ATAC (5kb merged) (right) modalities from the multiome dataset, colored by batch and cell type. (B) UMAP of the MVAE PoE latent representation for the multiome dataset, showing improved batch mixing and cell type separation. (C) UMAPs of the preprocessed RNA (left) and protein (right) data for the CITE-seq dataset, colored by batch and cell type. (D) UMAP of the MVAE PoE latent space for CITE-seq, demonstrating effective integration. Benchmarking results for the multiome dataset (E) and CITE-seq dataset (F), comparing bio-conservation and batch correction scores across methods. Dashed lines indicate average scores across all tested tools.

For the 10X multiome dataset, we applied minimal preprocessing: scRNA-seq data was log-normalized using Scanpy^36^, and scATAC-seq reads were merged into 5 kb genomic windows to reduce sparsity while retaining biological relevant information (see Methods). No further feature selection or dimensionality reduction was applied prior to model input. The scRNA-seq modality displayed moderate cell type separation but retained batch effects while the scATAC-seq modality showed poor cell type separation despite good batch mixing (Fig. 4A). After training MIMA, the PoE latent space showed batch mixing and preserved cell type identity, even revealing subpopulations in many cases (Fig. 4B). We next evaluated the model on a scCITE-seq dataset, which combines antibody-oligonucleotide conjugates with scRNAseq to simultaneously quantify the presence of specific proteins and the transcriptome of single cells. The scRNA-seq data was preprocessed as described in Methods, while the Antibody-Derived-Tags count matrix, quantifying the occurrence of specific proteins, was used as provided by the authors^34^. In the input data, some cell types were separated in the scRNA-seq, though batch effects were evident (Fig. 4C). The quantified proteins showed little structure in the data, but clusters were diffuse across the embedding and batches were poorly mixed. After training of MIMA, the integrated PoE space showed well-defined cell type clusters and strong batch correction (Fig. 4D). Notably, cell types previously dispersed across the embedding, such as CD14+ and CD16+ Mono, formed cohesive and well-separated clusters.

### Comparing MIMA’s performance against methods tailored for specific data modalities

We again benchmarked MIMA against state-of-the-art methods using the *scIB* metrics package (Fig. 4E-F). *Harmony* performed well on batch correction (especially for ATAC) but poorly on bio-conservation. Conversely, MOFA+ preserved biological information but struggled with batch correction. *MultiVI*^9^, a method specifically designed for single-cell multiome integration, achieved strong bio-conservation but average batch correction. Strikingly, MIMA’s biological (PoE) latent space outperformed all other methods in overall integration quality, obtaining an effective balance between correcting technical variability and preserving relevant biological signals. It surpassed *multiVI* in batch correction, despite the latter incorporating modelling assumptions specific for these particular data types, which might not hold for other assays or datasets.

Similarly for the scCITE-seq dataset, we benchmarked MIMA against other state-of-the-art methods, including *totalVI*^10^, which is specifically designed for integrating scCITE-seq data. More general methods like PCA, *Harmony*, and *Scanorama* underperformed, especially on the RNA modality (Fig. 4F). MOFA+ again showed above-average bio-conservation, but failed to adequately correct batch effects. While *totalVI* achieved the best overall score (0.58), MIMA was a close second (0.54), despite lacking specialized modeling assumptions for these specific data types.

Across all experiments (Figs. 2–4), our results demonstrate that MIMA is a powerful and generalizable framework for multimodal data integration and batch correction. In the context of spatial omics, it was able to learn biologically meaningful representations, correct strong batch effects (particularly in MSI), and reveal spatially coherent tissue structures (Figs. 2–3). Importantly, the model not only preserved expert-defined biological annotations but also revealed molecular gradients and potential pre-malignant regions. MIMA performed competitively across diverse omics types, without requiring task-specific reengineering, highlighting the power and adaptability of our general-purpose architecture.

## Discussion

The study of complex tissue biology often requires integrating multiple high-throughput assays that capture thousands of heterogeneous molecular features, alongside complementary modalities such as high-resolution microscopy. To extract meaningful insights, these high dimensional datasets must be projected into lower-dimensional representations that disentangle the sources of variation. This decomposition should contain components that reflect variability shared across modalities and samples, modality-specific signals present across multiple samples, and sample- or batch-specific variability. Several approaches have been proposed to tackle this integration problem, with only some of them also addressing the decomposition of the sources of variation. Additionally, most of these frameworks are tailored towards specific combinations of modalities. Here we propose a novel modular architecture that is modality-agnostic, can accommodate any number of modalities, and simultaneously disentangles the different sources of variation. We demonstrate its application in a variety of spatial and single-cell multi-omic datasets, including the first reported batch-corrected and integrated representation of spatial transcriptomics and lipidomics data. Our method successfully captured technical variability even in modalities prone to batch effects, like mass-spectrometry imaging data, that would otherwise prevent effective cross-sample studies. This advancement allows for the spatial co-localization of transcriptomic and metabolomic features in tissue, unlocking new insights into disease biology and tissue organization that were previously unattainable.

By integrating the different modalities our model is able to automatically predict expert human manual annotations with high accuracy and resolution. In contrast to manual annotations, MIMA captures continuous transitions between different states. MIMA detects molecular patterns consistent with early tumorigenic changes in regions labelled as non-tumour or pre-malignant by the pathologist. This suggests that our approach can reveal molecular alterations not yet visible by histological inspection alone, offering new avenues for precision diagnostics and early detection. Additionally, MIMA enables cross-modal translation, providing a mechanism to impute one modality from another. This is particularly valuable in settings where not all modalities may be available for every sample. Successful cross-modal translation also indicates that our shared representations capture meaningful relationships between different modalities.

In the prostate cancer dataset, the data acquisition process prioritized biological heterogeneity, resulting in a challenging but realistic setting where some annotations, such as rare tumor subtypes like ISUP 2 and 4, are present in only one sample each. This high inter-sample variability, combined with confounding batch effects, hinders the disentanglement of technical and biological signals. To reach fully robust cross-modal translations, more comprehensive datasets will be required that cover rare subtypes in multiple biological replicates. Alternatively, zero-shot or online learning approaches could be explored, however their accuracy remains to be seen in cross-modal translation. This represents a key challenge in real-world multi-sample omics datasets, where the scarcity of certain classes can limit the robustness of cross-modal translation, underscoring the need for integrative models like MIMA that leverage all available data modalities to improve biological inference under such constraints.

Another key feature of MIMA is that it is unsupervised and modality-agnostic. The model makes no assumptions about the types or number of input modalities, making it inherently scalable and adaptable to different experimental setups. This follows current technological trends where more modalities are measured on the same individual sample^1^. Specialized tools like *totalVI* or *multiVI* work well on specific data types (e.g., sc CITEseq, scRNAseq + scATACseq), but lack the flexibility to generalize to other datasets. By applying MIMA to two distinct single-cell datasets: multiome (RNA + ATAC) and CITE-seq (RNA + protein); we show that it performs similarly to state-of-the-art tools specifically tailored towards these technologies. The modular design of MIMA allows users to easily incorporate additional modalities without redesigning its architecture. Moreover, MIMA takes all input features as is, not relying on extensive prior preprocessing, such as dimensionality reduction and/or feature engineering. This allows the model to learn relationships between the actual molecular features directly from the data.

As with most deep learning approaches, interpretability remains a challenge: while MIMA successfully captures correlations between molecular features across modalities, it is unclear whether these relationships reflect true underlying biology. Applying explainability techniques such as integrated gradients^37^ or SHAP values^38^ could help understand the features driving the model’s predictions. However, most of the current explainable AI methods are not well-suited for highly correlative, high-dimensional omics data, underscoring the need to develop new interpretability approaches tailored to this domain. Incorporating perturbation datasets, where causal mechanisms can be more directly probed, could provide a powerful framework both to validate and to refine the biological relevance of the learned representations. Our architecture disentangles variation into shared and modality-specific components. Further improvement of the latent space disentanglement will require future methodological developments such as orthogonality between latent dimensions, enabling clearer attribution of signals to their biological or technical origin. Additionally, explicitly modelling more complex covariate structures (e.g., patient metadata, hierarchical batch effects or temporal dependencies) could also improve the learned representations. MIMA’s modular design allows for the pretraining of the encoder and decoder of each modality separately, exploiting the vast amounts of publicly available unimodal data. However, the role and impact of such pretraining on multi-modal data integration remains unclear^39^. These opportunities have the potential to improve MIMA’s accuracy and robustness, and also expand its utility as a foundation for the next generation of AI-based tools in computational pathology and multi-omics integration.

In summary, we present MIMA as a general-purpose, modular, and unsupervised framework for multi-omics data integration and batch correction. It enables robust biological signal extraction across spatial and single-cell datasets, accurately reflects known pathological annotations, and even reveals previously unrecognized disease states. MIMA matches state-of-the-art methods tailored to specific tasks, while remaining broadly applicable to a wide array of data modalities, including novel combinations, such as spatial transcriptomics and mass spectrometry imaging. This work provides a useful tool for current multi-omics studies and also serves as a foundation for next-generation AI systems in digital pathology, with the potential to enhance diagnosis, prognosis, and precision medicine by harnessing the full complexity of molecular tissue profiling.

## Materials and Methods

### Datasets

We evaluated our approach on three multimodal datasets, consisting of one spatial multi-omics dataset and two single-cell multi-omics datasets using different molecular modalities to illustrate the modality-agnostic nature of MIMA.

### Spatial Prostate Cancer (PCa) Dataset

This dataset is described in Zhang et al.^19^ and comprises spatially-resolved multi-omics data from eight prostate cancer (PCa) patients. For each patient, two tissue samples were collected: one from a tumorous region and one from a cancer-distal region, expected to be non-tumorous, enabling intra- and inter-patient comparisons. Each sample was profiled using three spatial modalities on consecutive tissue sections: spatial transcriptomics using the 10x Genomics Visium platform, Mass Spectrometry Imaging (MSI) for spatial lipidomics and a third modality derived from histological imaging. Morphological features were extracted from Hematoxylin and Eosin (H&E)-stained slides, from the tissue sections used for MSI, followed by feature extraction using a pre-trained convolutional neural network^25^, yielding 512-dimensional image embeddings per spatial spot. Altogether, the dataset includes 18950 genes, 12510 lipid m/z features, and 512-dimensional morphological embeddings across 42475 spatial spots from the 16 tissue samples (2 per patient).

Furthermore, the dataset includes detailed manual pathological annotations provided by expert pathologists specialized in prostate cancer. Annotated tissue types include: stroma, normal epithelium, atrophy, cribriform, necrosis, nerve, PIN (prostatic intraepithelial neoplasia), inflammatory infiltrate, other, tumor ISUP grades 2 through 5. Non-annotated spots were marked as “unlabelled”.

### Single-cell Multiome Dataset

The second dataset, obtained from the 2021 NeurIPS OpenProblems challenge^34,40^, contains single-cell multiome data comprising gene expression (scRNA-seq) and chromatin accessibility (scATAC-seq) from bone marrow mononuclear cells of 12 healthy donors. It includes a total of 69249 cells, with measurements for the expression of 13431 genes and 116490 chromatin accessibility ATAC-seq peaks.

To reduce the high sparsity and dimensionality of the ATAC modality, we preprocessed the ATAC peaks by aggregating them into 5 kb genomic bins. This resulted in a less sparse and more informative feature space of 40110 ATAC aggregated bins. The RNA modality was preprocessed with a basic total count normalization and log transformation with *Scanpy*^36^ from the count data.

### Single-cell CITE-seq Dataset

The third dataset also originates from the 2021 NeurIPS OpenProblems challenge^34,40^, containing CITE-seq data with paired gene expression and surface protein abundance measurements. The data consists of 90261 single cells from bone marrow mononuclear cells of 12 healthy human donors, assayed for 13953 genes and 134 surface proteins. The RNA modality was preprocessed the same way as in the multiome case. The proteomics modality was used as provided by the authors.

### Classification of Pathologist Annotations

To evaluate the power of the learned latent spaces to predict manual expert annotations, we trained an ensemble of random forest classifiers (ensemble classifier, EC), implemented using the BalancedBaggingClassifier (using a Random Forest Classifier as base estimator) from the imbalanced-learn library (*v0*.*13*.*0*)^41^, to mitigate the strong class imbalance present in the dataset (Supplementary Fig. 1C). For this classification task, six annotation classes were excluded due to their low representation in the dataset or lack of clear biomedical meaning: nerve, necrosis, tumour ISUP 2, tumour ISUP 4, other and unlabelled.

A binary one-vs-rest classifier was trained for each of the remaining annotation classes: atrophy, cribriform, inflammatory infiltrate, normal epithelium, PIN, stroma, tumor ISUP 3 and tumor ISUP 5. Classification performance was assessed using accuracy, precision, recall, and F1-score, computed only when the test set contained ≥3 instances of the target annotation. A global performance metric was then obtained by averaging the four metrics across all binary classifiers.

To avoid sample-wise overfitting, we used leave-one-out cross-validation (LOOCV) across the 16 PCa samples, training the EC on 15 samples and testing on the held-out sample in each fold. Metrics were averaged across folds to produce a global score.

As in some cases certain annotations are exclusive to individual samples, and therefore LOOCV would prevent the EC from learning to predict these rare labels, we also performed a different within-sample 4-fold cross-validation (4CV) scheme. In 4CV, we randomly partitioned sample datapoints so that 25% of the spots were used as the test set and the remaining 75% as the training set, ensuring all annotation types were represented during training. This process was repeated four times, allowing full prediction coverage while preserving label diversity and minimizing biases.

### 3D-RGB UMAP

To visually interpret the structure of the learned latent space, we compute a 3D UMAP embedding from the latent representations. Each of the 3 UMAP axes is min-max scaled to the [0, 1] range and then mapped to the red, green, and blue (RGB) color channels, respectively. This RGB encoding allows each spatial spot to be assigned a unique color based on its position in the 3D embedding space. As a result, spatially or biologically similar spots appear in similar colors, forming smooth gradients, while dissimilar spots are easily distinguishable by contrasting colors. This provides an intuitive and information-rich visualization of the latent space organization and the relationships between spatial spots.

### Highly variable features selection

To reduce the impact of dropouts (Supplementary Fig. 1A) and to improve the robustness of downstream analyses, we perform feature selection by identifying the top 1000 highly variable features (HVFs) for the RNA and MSI data. First, we compute the variance of each feature across all samples and retain the top 2000 most variable features. From this subset, we further select the 1000 features with the highest mean expression or abundance, ensuring that selected features are both variable and consistently expressed.

### Gene enrichment score

To assess the molecular consistency of the model predictions with known prostate cancer biology, we computed enrichment scores based on a curated set of 578 genes previously associated with prostate cancer^27^. Enrichment scores were calculated using the *scanpy*.*tl*.*score_genes* function (Scanpy v1.11.1), which computes, for each spot, the average expression of the selected gene set subtracted by that of a reference set of genes matched for expression distribution. This generates a continuous enrichment score reflecting the relative activation of the prostate cancer gene signature at single-spot resolution.

### Quantitative Benchmarking of State-of-the-Art Integration Tools

To systematically evaluate the quality of our model, we benchmarked the learned latent representations against those of several widely used batch correction and data integration tools using the *scIB* metrics package (*v0*.*5*.*3*)^29^. This benchmarking framework computes a total of 12 quantitative metrics: 7 of these metrics assess biological conservation (i.e., how well the embedding retains biologically meaningful structures based on ground truth labels), and 5 evaluate batch correction (i.e., how well technical variation is removed) (Supplementary Table 1). A composite score is then derived by averaging the bio-conservation and batch correction metrics, offering a balanced evaluation of each method’s performance according to these two different aspects.

To ensure fair comparisons, all methods that allowed customization of the latent dimensionality were configured to produce 100-dimensional embeddings, consistent with our MIMA architecture. Tools that failed to run on our datasets (e.g., LIGER^42^ and SpatialGlue^33^) were excluded from the analysis. SpatialGlue, in particular, was incompatible due to its requirement for specific input modalities.

We compared MIMA to the following tools:

- *MOFA+*^11^: A statistical factor analysis framework for modelling multi-omics data using a linear generative model trained via variational inference. MOFA+ infers latent factors that capture the shared and modality-specific sources of variation.
- *scVI*^32^: A deep generative model for single-cell RNA-seq data using variational autoencoders. It serves as the foundation of the scvi-tools ecosystem, which includes several multimodal extensions.
- *TotalVI*^*10*^ and MultiVI^9^: Extensions of *scVI* tailored for the integration of single cell gene expression data with proteomics data(scRNAseq + scCITEseq) and multiome data (scRNA + scATACseq) respectively, designed to jointly model paired modalities and correct for batch effects.
- *Harmony*^30^: An algorithm that projects single-cell data into a shared space while aligning data across batches, typically used following PCA preprocessing.
- *Scanorama*^31^: A method for batch correction across scRNA-seq datasets that detects and merges shared cell populations across batches via manifold alignment.
- Principal Component Analysis (PCA): A classical linear method used as a baseline for dimensionality reduction and often as input for integrative models like *SpatialGlue*. We computed PCA on each modality individually and on the “unintegrated” case, where all modalities were concatenated before PCA was applied.

Additionally, to further assess the integrative performance of the MIMA architecture, we compared it to a unimodal setting (i.e. MIMA-Unimodal). For this, we trained a modality-specific variant of our architecture removing the multimodal integration step to serve as a unimodal baseline. In this configuration, only the modality-specific and batch latent spaces were updated, and the PoE (shared latent space) was not used.

### Model training

MIMA is trained using a composite loss function that disentangles shared and modality-specific (private) objectives, enabling modality-agnostic integration and cross-modal translation in an unsupervised fashion.

### Shared loss

The integrated (PoE) latent 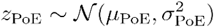 space is derived from the shared latent space of each individual modality *m* 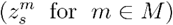 using a Product-of-Experts (PoE) formulation^18^ across all input modalities. The shared spaces capture shared biological signals and are used both for reconstructing the original input from their PoE combination and for performing cross-modal translation.

The shared loss consists of:

- Reconstruction Loss (PoE): For each modality *m* ∈*M*, let *x*_*m*_ be the input and 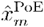 its reconstruction using the PoE latent space. The reconstruction error is measured using Mean Squared Error (MSE):

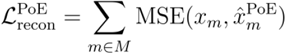
- Translation Loss: Each modality is translated to every other using its shared latent representation *z*_*s*_, producing predicted outputs 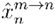 for *n*∈*M, n* ≠*m*. The translation loss is computed as:

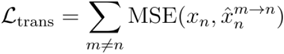
- KL Divergence (PoE): The PoE latent space z_PoE_ is regularized towards a standard Gaussian prior by minimizing the KL divergence:

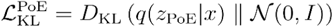

The total shared loss becomes:

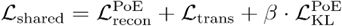

### Private loss

Modality-specific batch-specific variation is also encoded into private latent spaces 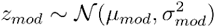 and 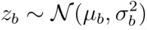, capturing information specific to that modality or sample, respectively.

The private loss is composed of:

- Reconstruction Loss (Private): The modality is reconstructed from its private components 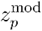

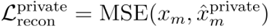
- KL Divergence (private):

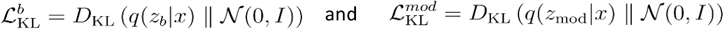

The shared space for each modality 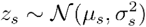 is also regularised towards as a standard Gaussian in the private loss component:

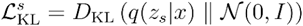

The total private loss for each modality becomes:

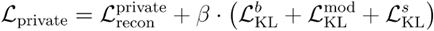

### Total loss

The final loss used to train the model is the sum of the shared and private losses:

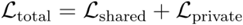

### MIMA parameters and hyperparameters

All training hyperparameters were kept consistent across modalities and experiments, and were selected after thorough manual hyperparameter tuning to balance performance and generalization. A dropout rate of 0.1 was applied in the encoder/decoder layers and 0.3 in the latent space layers to reduce overfitting, models were trained for 50 epochs using the Adam optimizer^43^ with a learning rate of 1e-4, a batch size of 128, 80-20 train-test split, ReLU activation, and a KL regularization coefficient (β) of 0.01 for all submodules and PoE regularizations, except for the CITE-seq data, where the PoE β was set to 0.1 and the RNA submodule to 0.001. For the PCa data, the MIMA model was composed of three modality-specific submodules: RNA, MSI, and morphology. RNA and MSI encoders each consisted of two hidden layers with 300 units, projecting the input into a 100-dimensional latent space. For the morphology modality, due to its lower input dimensionality (512 features), the input was directly projected into the latent space without encoder layers. The decoders for all three modalities mirrored their respective encoder architectures to reconstruct the original input space.

For the multiome data, the RNA and ATAC submodules shared the same architecture used for the PCa RNA and MSI encoders: two hidden layers of 300 units followed by projection into a 100-dimensional latent space, with decoder symmetry. Dropout rates, learning rate, batch size, activation function, and training schedule were kept identical to the PCa setup. This consistency allows for direct comparability across datasets and ensures robustness of the modular design.

For the CITE-seq data, the RNA submodule was kept the same as in the PCa and multiome configurations. The protein modality was handled equally to the morphology data in the PCa dataset: no hidden layers were used between the input and latent space due to its low dimensionality. The latent dimensionality was again set to 100.

## Data availability

The datasets used for this study, together with the trained model parameters for each dataset for figure reproducibility, can be found in Zenodo: 10.5281/zenodo.17369670.

## Code availability

The source code for this project can be found in our github: https://github.com/sifrimlab/MIMA.

## Author contributions

JIAL: conceptualization and implementation of the model. Applied the model to the described datasets, analysis of the datasets, visualization, writing and editing of the manuscript. GP: conceptualization and implementation of the model. Writing and editing of the manuscript. LV: refined codebase, wrote documentation and created reproducible environments. WZ: PCa data analysis and creation of morphological embeddings. XS: PCa data generation. SV: PCa data analysis. SK: PCa data generation. KV: PCa data generation plus its supervision. DW: PCa data analysis. AI: applied and tested model with the datasets. RS: applied and tested model with the datasets. JI: PCa data generation. TS: PCa data generation. SRE: PCa data generation. ML: PCa data generation and interpretation. FB: PCa data generation. TG: PCa data generation. SJ: PCa data generation. MC: PCa data analysis. NV: PCa data analysis. TV: PCa data generation plus its supervision. JS: PCa data generation plus its supervision. JJ: supervised the work, writing and editing of the manuscript. AS: supervised the work, writing and editing of the manuscript.

## Acknowledgments

JIAL was supported by a PhD Fellowship from the Research Foundation Flanders (FWO) strategic basic research (1SF1522N and 1SF1524N).

LV is supported by the European Union’s Horizon 2020 research and innovation programme under the Marie Skłodowska-Curie Actions Doctoral Network ‘PROSTAMET’, Grant Agreement No. 101120283 (HORIZON-MSCA-2022-DN-01)

WZ was supported by a Baekeland PhD grant from Flanders Innovation and Entrepreneurship (VLAIO) - HBC.20192204.

XS was a recipient of a FWO PhD fellowship (SB/1S49218N) and was in part supported by Opening the Future (OtF).

SRE acknowledges support from the Australian Research Council (FT190100082).

TS acknowledges support through the Australian Government Research Training Program Scholarship.

SV was sponsored by an FWO PhD Fellowship strategic basic research (1S93320N).

ML was supported by grants: DoD HT9425-25-1-0385 and P01 CA265768 (NCI).

KV, SK and TV were supported by KU Leuven (IDN/19/039, Opening the Future (OtF), Leuven Future Fund (LFF)).

This work was supported by a grant from the Belgian Foundation Against Cancer (STK), an EU HORIZON-MSCA-DN project 101120283 (PROSTAMET), a Focus Group grant from the LKI Fund for Innovative Cancer Research (FIKO), the KU Leuven Opening the Future campaign, KU Leuven Incubation financing Lipometrix, a grant PRISMO from the Flemish Resilience Plan, FWOEOS projects 30837538 (DECODE), G0F6718N (SeLMA), FWO-SBO projects S001623N (LIPOMACS), S005319N and S000825N (GLIOSELECT). Figure 2A was created with BioRender.com.

---

**Supplementary Figure 1.**
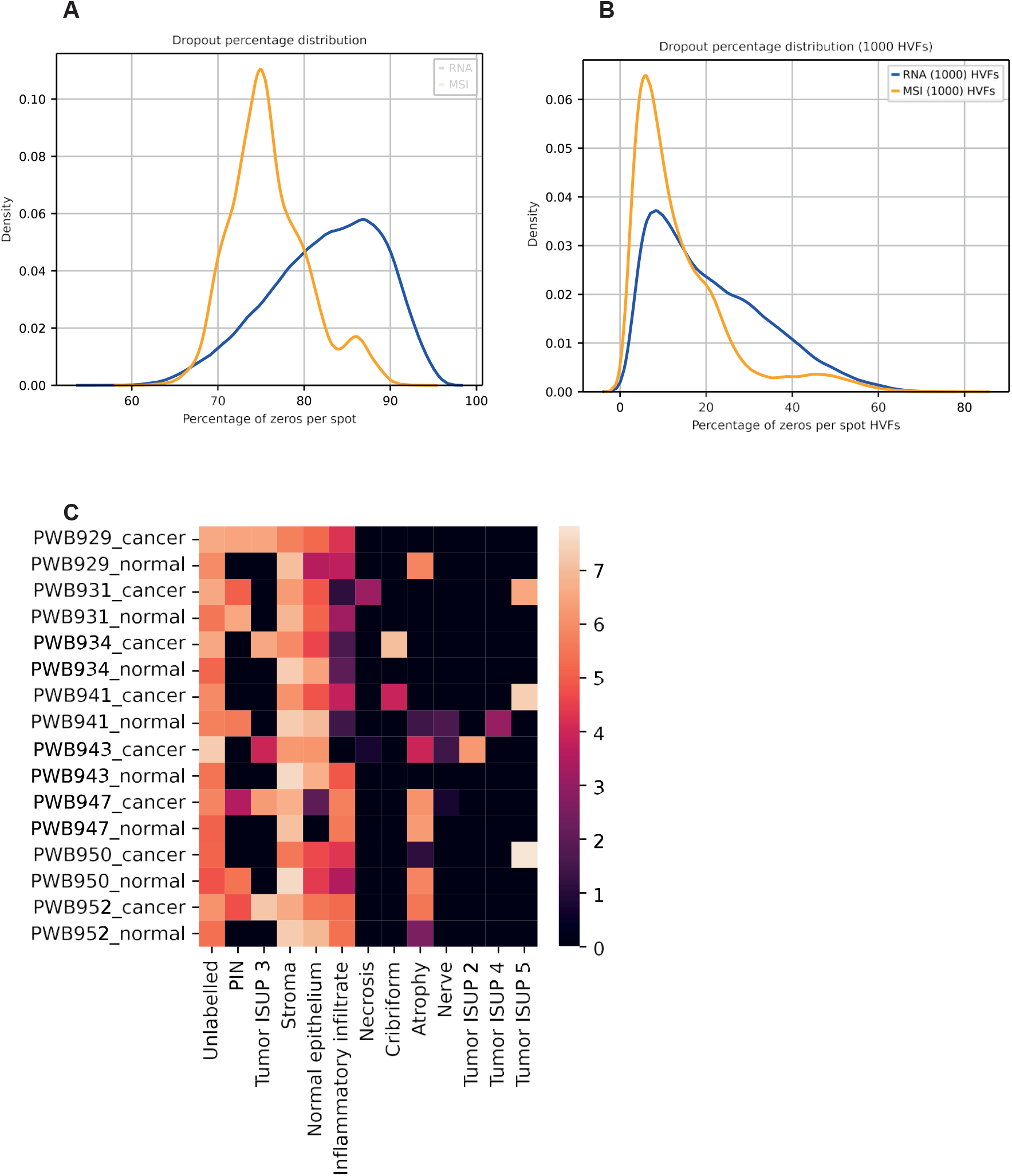
Prostate cancer (PCa) description. Distribution of the percentage of dropouts present in the dataset per spot for the whole dataset (A) and for the 1000 selected Highly Variable Features (HVFs) (B), for the RNA and MSI modalities. (C) Annotation distribution per sample, total counts were log-transformed for visualization.

**Supplementary Figure 2.**
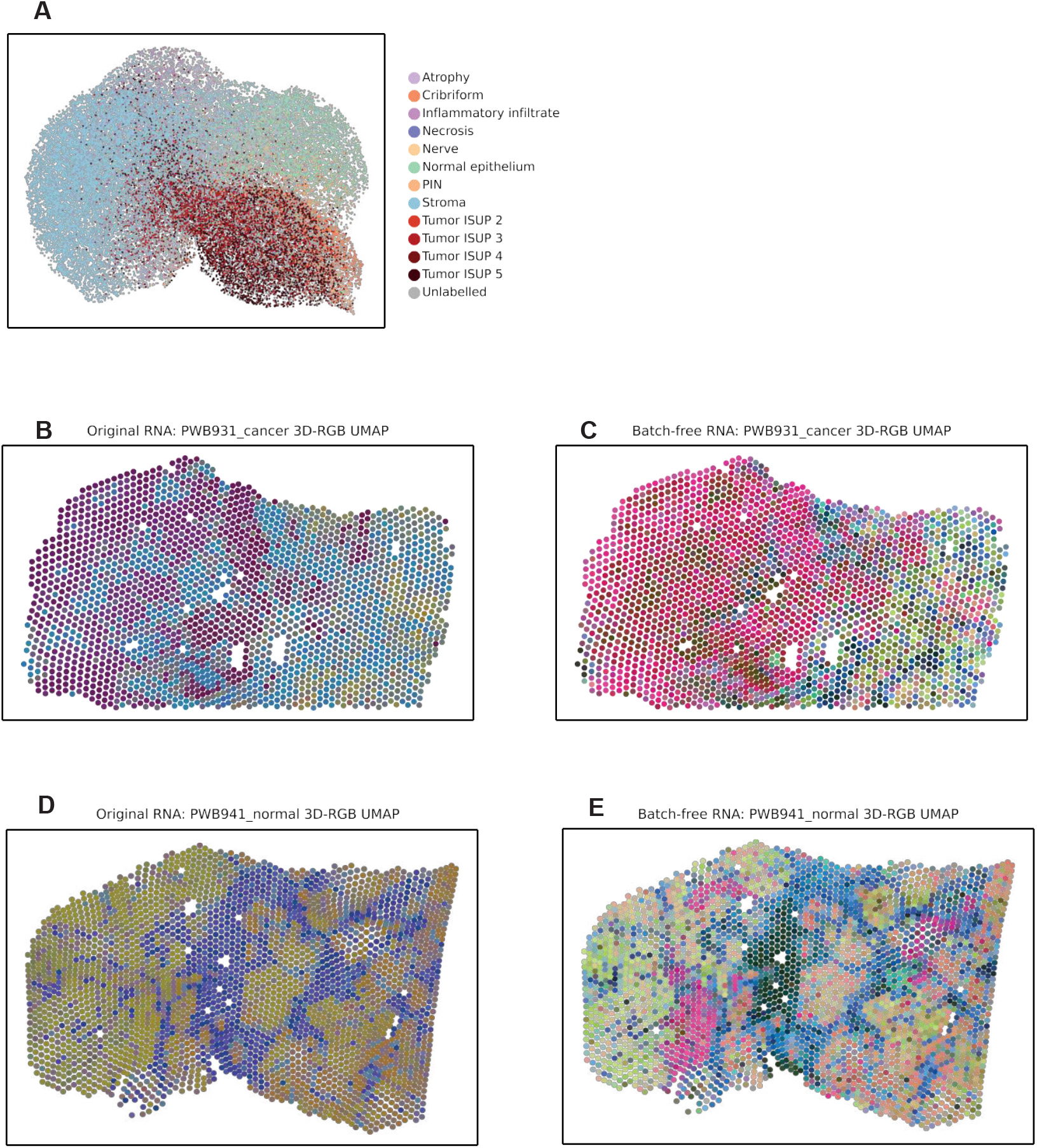
Batch correction performance of MIMA on the RNA modality. (A) UMAP embeddings of MIMA’s batch-corrected data for the RNA modality, colored by pathologist annotations. (B–E) 3D-RGB UMAP visualizations mapped onto tissue sections for samples PWB931_cancer and PWB941_normal, showing input (uncorrected) RNA data and MIMA batch-corrected outputs.

**Supplementary Figure 3.**
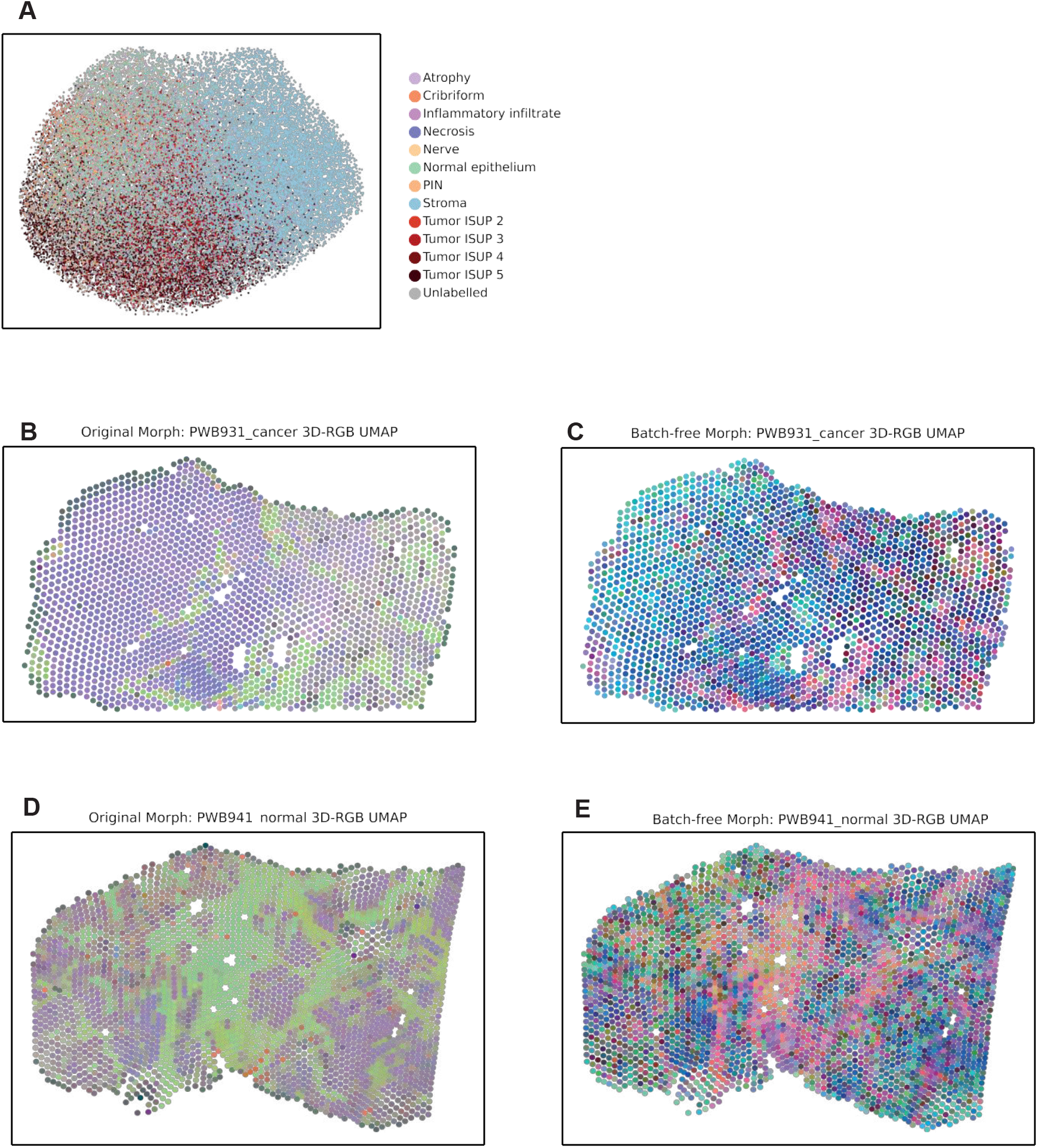
Batch correction performance of MIMA on the Morphology modality. (A) UMAP embeddings of MIMA’s batch-corrected data for the Morphology modality, colored by pathologist annotations. (B–E) 3D-RGB UMAP visualizations mapped onto tissue sections for samples PWB931_cancer and PWB941_normal, showing input (uncorrected) morphology data and MIMA batch-corrected outputs.

**Supplementary Figure 4.**
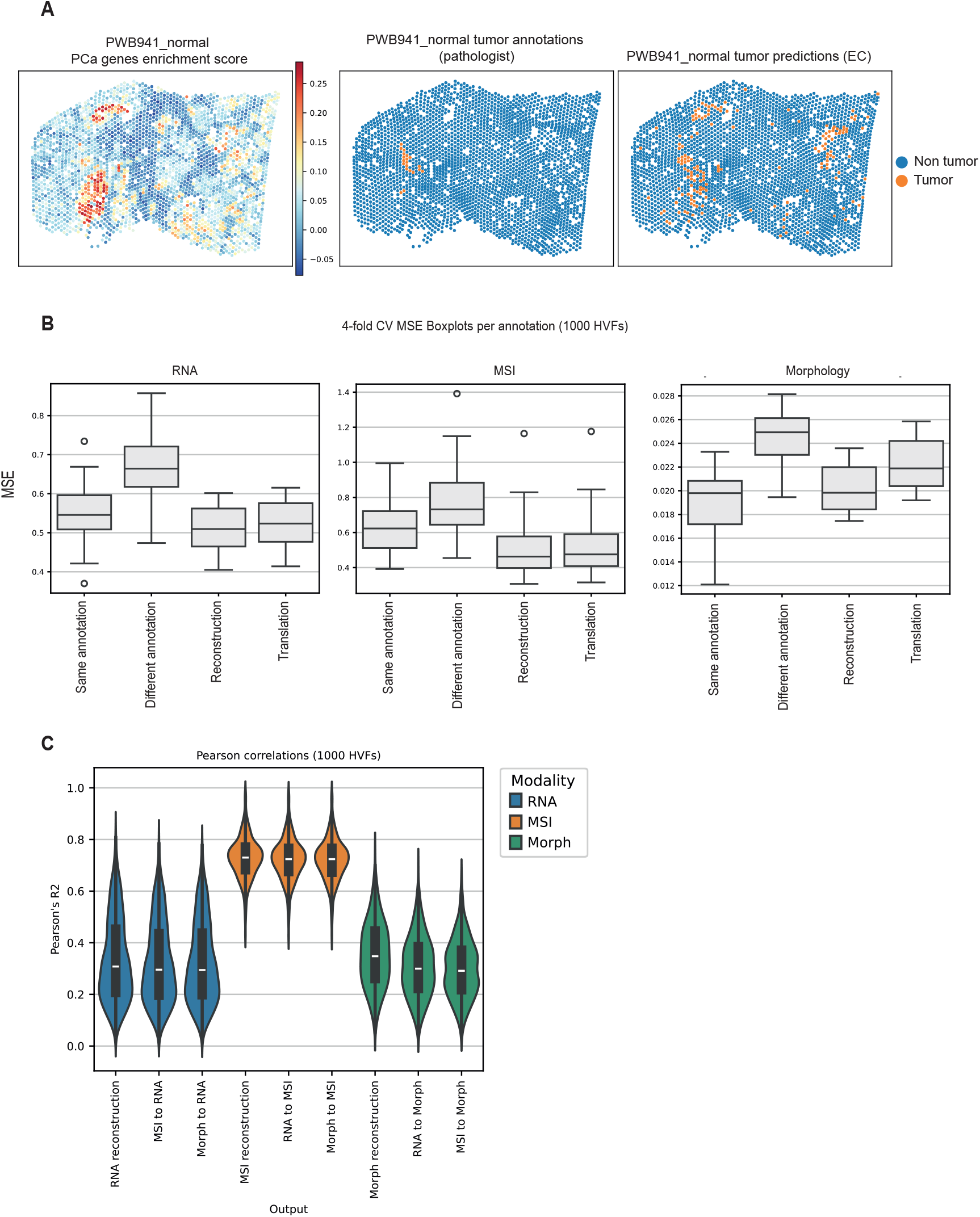
Validation of molecular predictions and model performance across modalities. (A) Spatial visualization of tumor detection for sample PWB941_normal, showing (left) enrichment scores of 578 prostate cancer–associated genes, (middle) tumor vs. non-tumor pathologist annotations and (right) predictions from the ensemble classifier. (B) Four-fold cross-validation (4CV) boxplots of mean squared error (MSE) across pathologist annotations for the top 1000 highly variable features (HVFs) in RNA, MSI, and Morphology modalities. (C) Pearson correlations between MIMA’s outputs (reconstructions and cross-modal translations) and input data for the 1000 HVFs.

