## Supplementary Figures for "Multi-omics integration and batch correction using a modality-agnostic deep learning framework"

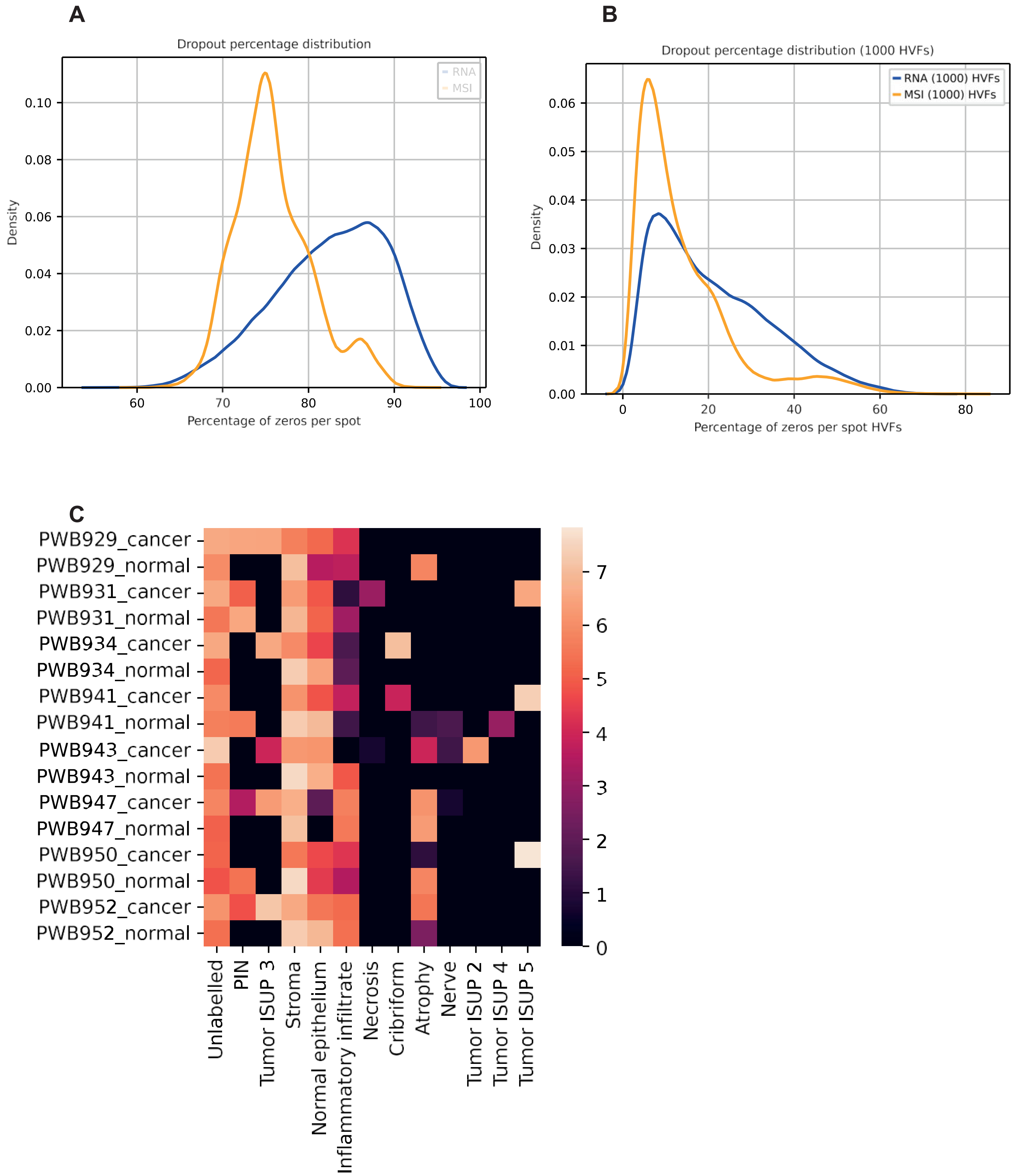


**Supplementary Figure 1. Prostate cancer (PCa) description.**

Distribution of the percentage of dropouts present in the dataset per spot for the whole dataset (A) and for the 1000 selected Highly Variable Features (HVFs) (B), for the RNA and MSI modalities.

(C) Annotation distribution per sample, total counts were log-transformed for visualization.

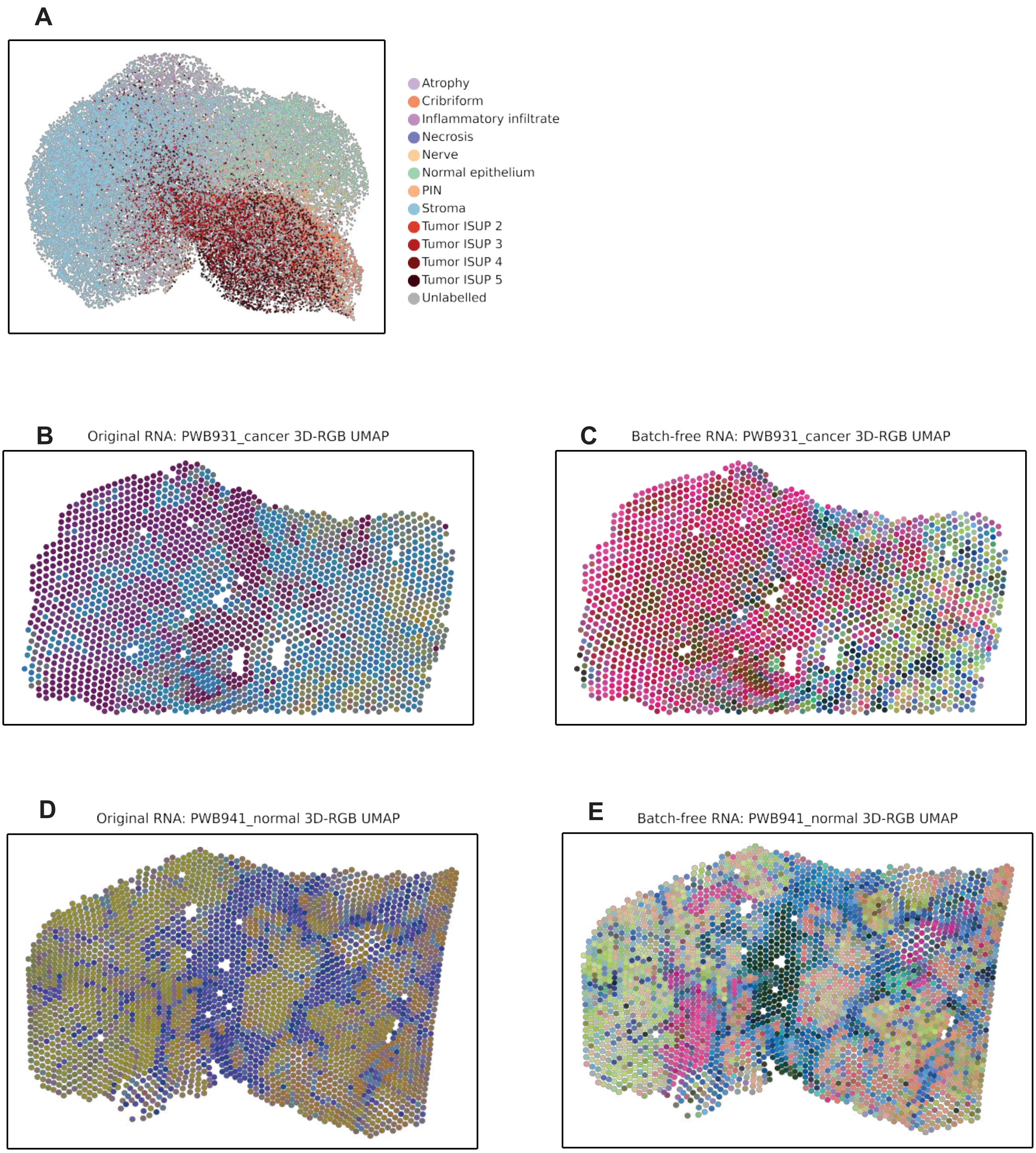


**Supplementary Figure 2. Batch correction performance of MIMA on the RNA modality.**
(A) UMAP embeddings of MIMA's batch-corrected data for the RNA modality, colored by pathologist annotations.
(B–E) 3D-RGB UMAP visualizations mapped onto tissue sections for samples PWB931_cancer and PWB941_normal, showing input (uncorrected) RNA data and MIMA batch-corrected outputs.

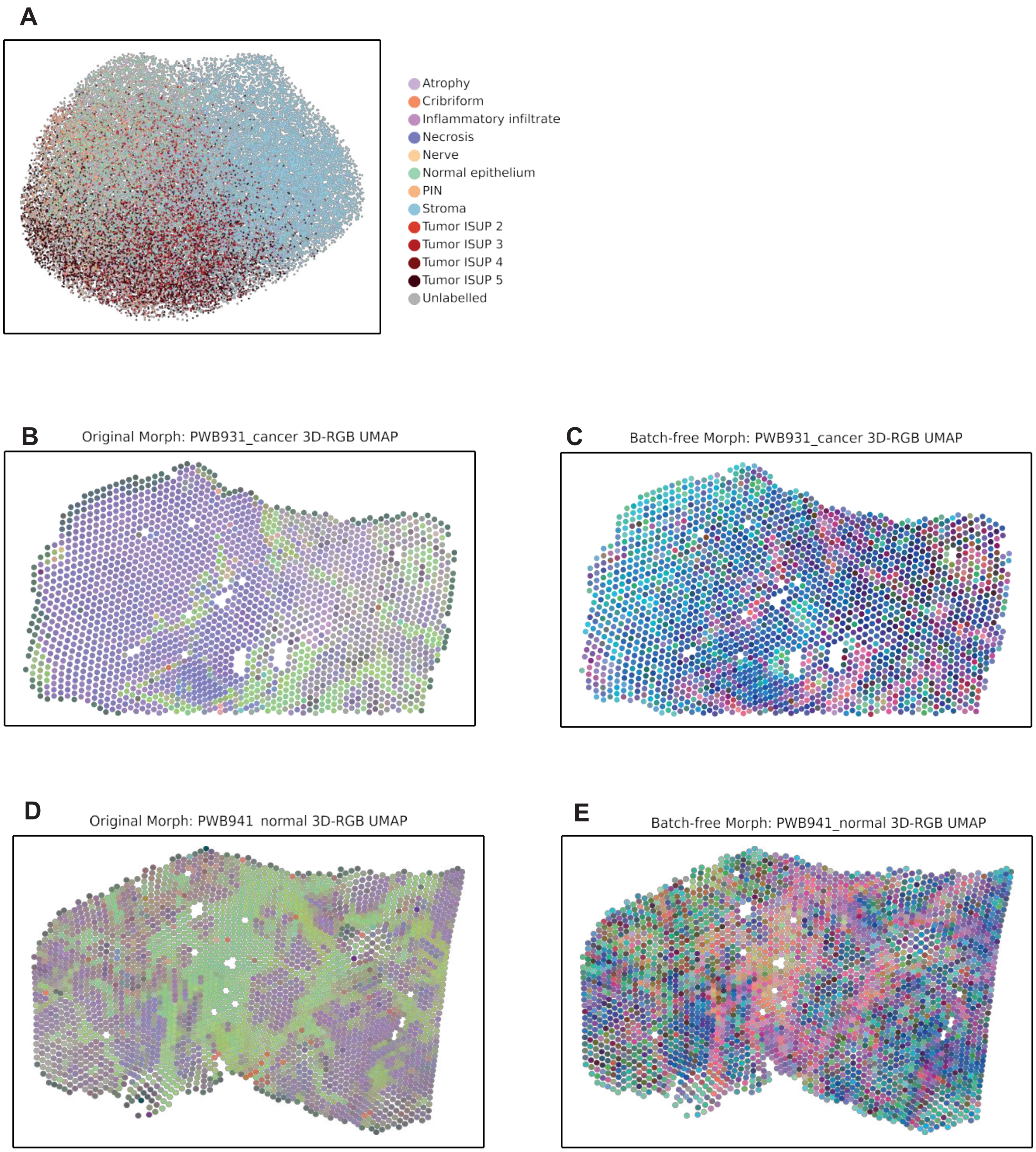


**Supplementary Figure 3. Batch correction performance of MIMA on the Morphology modality.**
(A) UMAP embeddings of MIMA's batch-corrected data for the Morphology modality, colored by pathologist annotations.
(B–E) 3D-RGB UMAP visualizations mapped onto tissue sections for samples PWB931_cancer and PWB941_normal, showing input (uncorrected) morphology data and MIMA batch-corrected outputs.

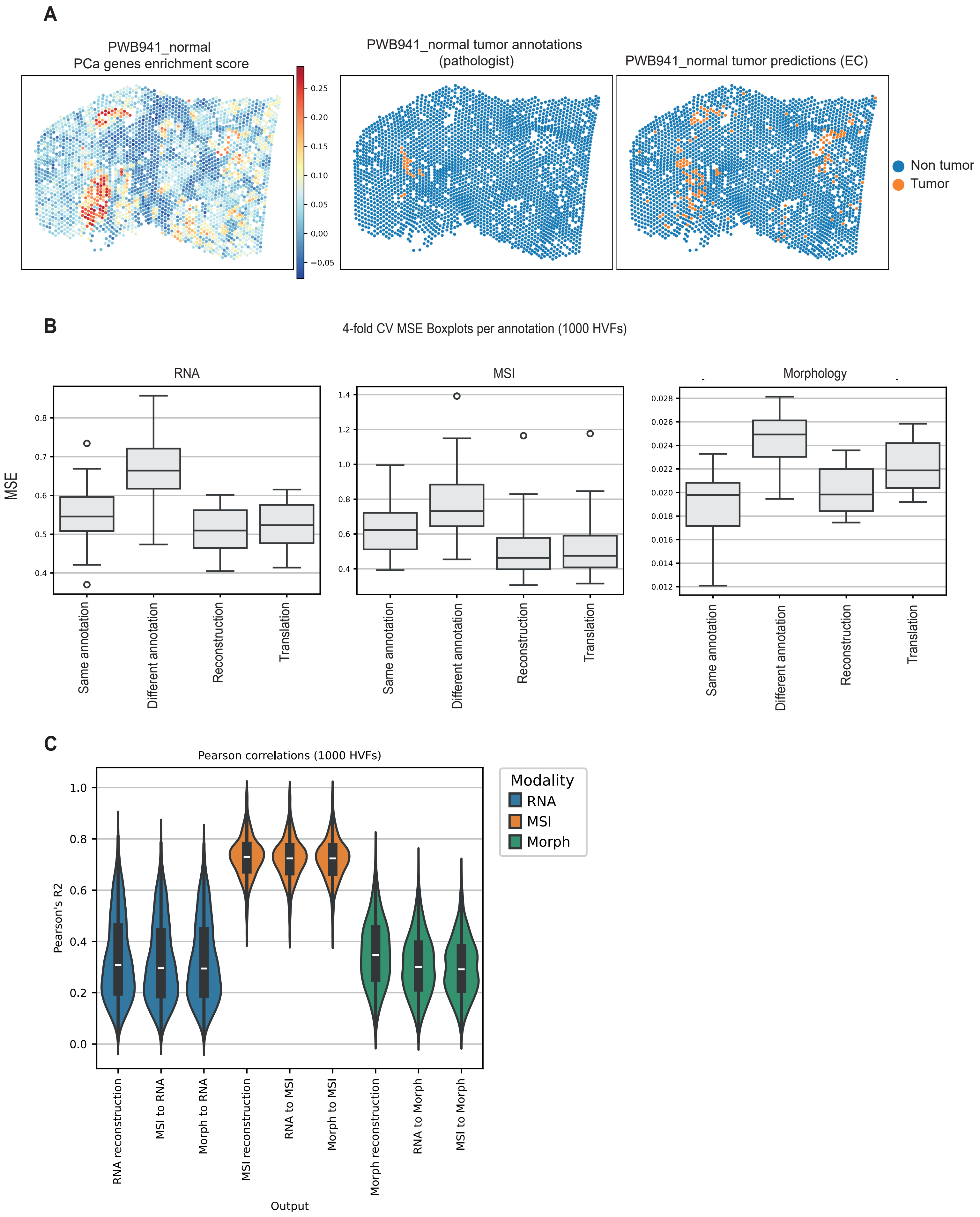
A
PWB941_normal
PCa genes enrichment score
0.25
0.20
0.15
0.10
0.05
0.00
−0.05
PWB941_normal tumor annotations
(pathologist)
PWB941_normal tumor predictions (EC)
Non tumor
Tumor
B
4-fold CV MSE Boxplots per annotation (1000 HVFs)
RNA
MSI
Morphology
MSE
0.8
0.7
0.6
0.5
0.4
1.4
1.2
1.0
0.8
0.6
0.4
0.028
0.026
0.024
0.022
0.020
0.018
0.016
0.014
0.012
Same annotation
Different annotation
Reconstruction
Translation
C
Pearson correlations (1000 HVFs)
Pearson's R2
1.0
0.8
0.6
0.4
0.2
0.0
Modality
RNA
MSI
Morph
RNA reconstruction
MSI to RNA
Morph to RNA
MSI reconstruction
RNA to MSI
Morph to MSI
Morph reconstruction
RNA to Morph
MSI to Morph
Output

**Supplementary Figure 4. Validation of molecular predictions and model performance across modalities.**

(A) Spatial visualization of tumor detection for sample PWB941_normal, showing (left) enrichment scores of 578 prostate cancer–associated genes, (middle) tumor vs. non-tumor pathologist annotations and (right) predictions from the ensemble classifier.
(B) Four-fold cross-validation (4CV) boxplots of mean squared error (MSE) across pathologist annotations for the top 1000 highly variable features (HVFs) in RNA, MSI, and Morphology modalities.
(C) Pearson correlations between MIMA's outputs (reconstructions and cross-modal translations) and input data for the 1000 HVFs.
